# High throughput photogrammetric measurement of morphological traits in free-ranging phototropic insects

**DOI:** 10.1101/699454

**Authors:** Mansi Mungee, Ramana Athreya

## Abstract

1. Remote measurement of morphological traits in free-ranging animals is very useful for many studies, but such non-invasive photogrammetric methods are limited to large mammals and have yet to be successfully applied to insects which dominate terrestrial ecosystem diversity and dynamics. Currently, insect traits are measured using collected specimens; the process of collection and maintenance of specimens can impose a heavy and unnecessary cost when such specimens themselves are not needed for the study.
2. We propose a rapid, simple, accurate, and semi-automated method for high-throughput morphometric measurements of phototropic insects from shape and size calibrated digital images without having to collect a specimen. The method only requires inexpensive, off-the-shelf, consumer equipment and freely available programming (*R*) and image processing (*ImageMagick*) tools.
3. We demonstrate the efficacy of the method using a data set of 3675 images of free-ranging hawkmoths (Lepidoptera: *Sphingidae*) attracted to a light screen. Comparison of trait values from a subset of these images with direct measurements of specimens using a scale showed a high degree of correspondence. We have also identified several error metrics which help in assessing the method in an objective manner.
4. Although this method was developed for nocturnal phototropic insects, it can be used for any other (small) animal that can be imaged on a simple graph paper. While this technique will be generally useful for a variety of studies of insect traits, we suggest that it is particularly suited as a commensal on multi-epoch and multi-location population monitoring of insects in the context of climate and land-use change, where repeated sampling obviates the necessity of collecting specimen every time. It will help in accumulating a large amount of reliable trait data on hundreds of thousands of individual insects without an overwhelming expenditure on collection, handling, and maintenance of specimens.

## Introduction

Morphological traits of insects, particularly body and wing sizes, have been key ingredients in a variety of studies spanning physiology (e.g. thermoregulation; Parmesan & Yohe, 2003; Sheridan & Bickford, 2011), macroecology (e.g. Blackburn & Gaston, 1994; Gillooly et al., 2001), ecogeography (e.g. Shelomi, 2012; Vinarski, 2014) and allometry (e.g. Voje et al., 2014). The need for standardized measures of morphological traits of insects and for large databases has been highlighted by many ecologists (e.g. Chown & Gaston, 2010; Moretti et al., 2017). Morphological measurements are traditionally carried out (only) on collected specimens using calipers or calibrated images from expensive microscopes. Wing measurements require proper mounting/spreading of the specimen (e.g. Moretti et al. 2017). Their measurement for a large number of individuals is the rate-limiting procedure for many investigations.

Photogrammetry is the method of estimating (free-ranging) subject length attributes such as size or distances from photographs (Baker, 1960). Typically, photogrammetry involves elaborate procedures and expensive equipment (e.g. Mahendiran, Parthiban, Azeez & Nagarajan, 2018) and so has been predominantly used for large taxa (mostly mammals, some birds) with small population sizes and high conservation priority (e.g. Durban & Parsons, 2006; Berger, 2012; Kurita, Suzumura, Kanchi & Hamada, 2012).

We have not come across photogrammetry of free-ranging phototropic insects, which are usually skittish and difficult to get to pose at the desired location. The usual practice is to collect all the individuals visiting a light trap. Apart from the challenges of actual measurement, we estimated that the cost of transforming a collected insect into a museum specimen *with long-term utility* costs 1-3 USD even in an inexpensive country like India. This can add substantially to project costs for taxa which arrive at a trap in their (tens of) thousands. Furthermore, the target taxon may be only a small fraction of the total individuals at the trap; the expense of processing unwanted but unavoidable specimens may surpass that for the targets.

We describe here a rapid procedure of photogrammetry to measure morphological traits of free-ranging phototropic insects without having to collect them. It can be used to measure the lengths of any visible morphological feature including body, wing, antennae, head, abdominal segments, elytra, etc, provided that it is being held parallel to the screen. The use of a screen as a substrate compels most individuals of a particular taxon (e.g. moths) into a similar posture, resulting in the uniformity necessary for automated processing. The procedure requires inexpensive, off-the-shelf, consumer equipment, and very little training and care while imaging the animals. We will also describe heuristics for quantifying errors of measurement and image quality (and hence usability).

We also suggest a different perspective of the generality of this method, ironically, arising from its particular suitability for moths. Moths are among the most abundant and diverse taxa and one of the most important herbivores and prey species in many ecosystems (New, 1997). They are easy to attract to light screens in large numbers. As insects go, they are relatively easy to identify down to genus from images, and to a lesser extent even to species. A procedure to collect body and wing sizes of large numbers of moths, as a commensal of population monitoring studies while not requiring a large amount of resources for preserving the specimens in a museum, would contribute to a global database and a variety of studies.

## Method & Materials

### A. Field Data

This procedure was developed for a study of traits of hawkmoths (*Lepidoptera*: Family *Sphingidae*) in Eaglenest wildlife sanctuary in Arunachal Pradesh, North-east India (Athreya, 2006; Mungee, 2018).

We attracted moths to a light screen made of a rectangle frame of fabric (1.6 x 1.1 m^2^) hanging from a portable and light-weight tetrapod (Figure 1a). An ultraviolet actinic lamp (8W T4-BL40) to attract moths and an optical lamp (8W CFC tube) to assist human vision were placed along the upper bar of the tetrapod for maximal effect. The lamps were powered by a 12V lead-acid battery through a DC-to-AC converter. The screen material (Figure 1c) was ordinary shirt fabric with thin checks (rectangular 7.0 x 7.3 mm^2^, for size and shape reference) on a white background (for maximum reflectance), stretched taut across the frame using elastic bands. The entire set up costs less than 250 USD in India at current prices.

**Figure 1.**
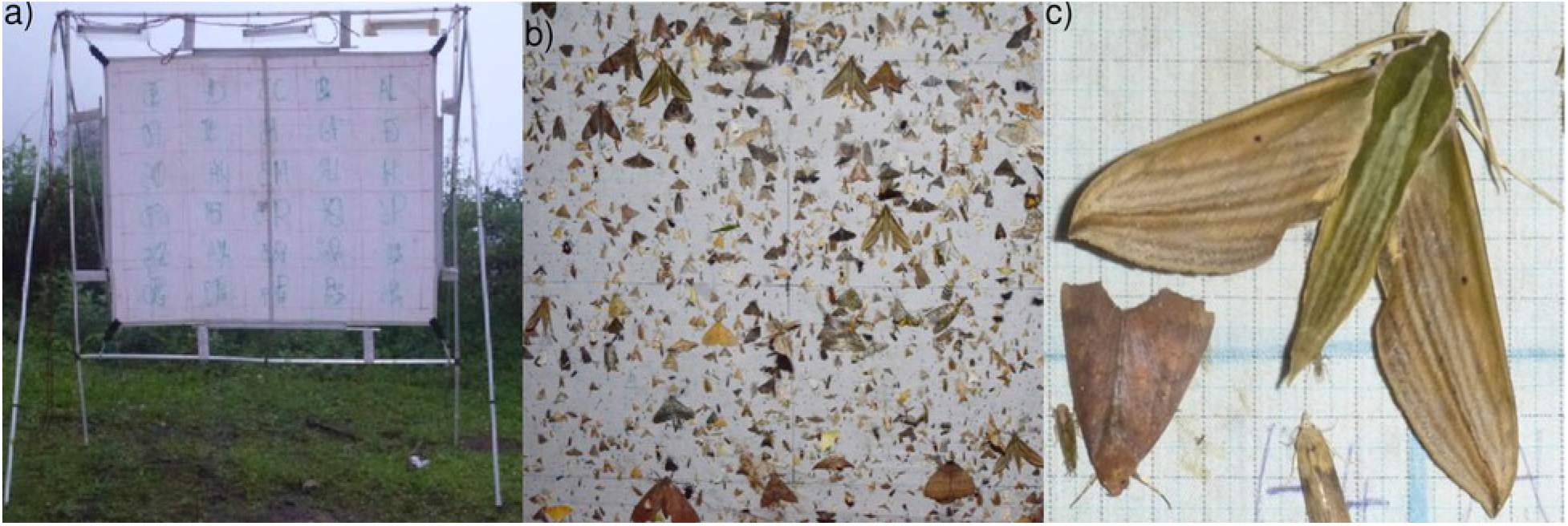
**a)** The portable moth-screen used to attract moths. **b)** Thousands of moth freely resting on the screen on a “good” night, **c)** A close-up of a Sphingid moth.

The number of moths and other insects varied from 5 to 3000 individuals each night (Figure 1b), depending on location, lunar phase, weather conditions, etc. We set up 2-8 screens at locations separated by 200-1000 m in elevation and 2-30 km along a forest road. The insects were photographed on the screen with cameras ranging from digital SLRs (Nikon D90 + 105 mm macro) to point-and-shoot (Panasonic Lumix DMC-TZ30).

These screens were manned by a large team of field staff with different levels of technical proficiency and educational backgrounds, resulting in images of variable quality. Most images suffer from a variety of distortions to a greater or lesser degree (Figure 2). We used the rectangular grid on the checked screen as the reference to post-facto calibrate the images for shape and size (Figure 3).

**Figure 2:**
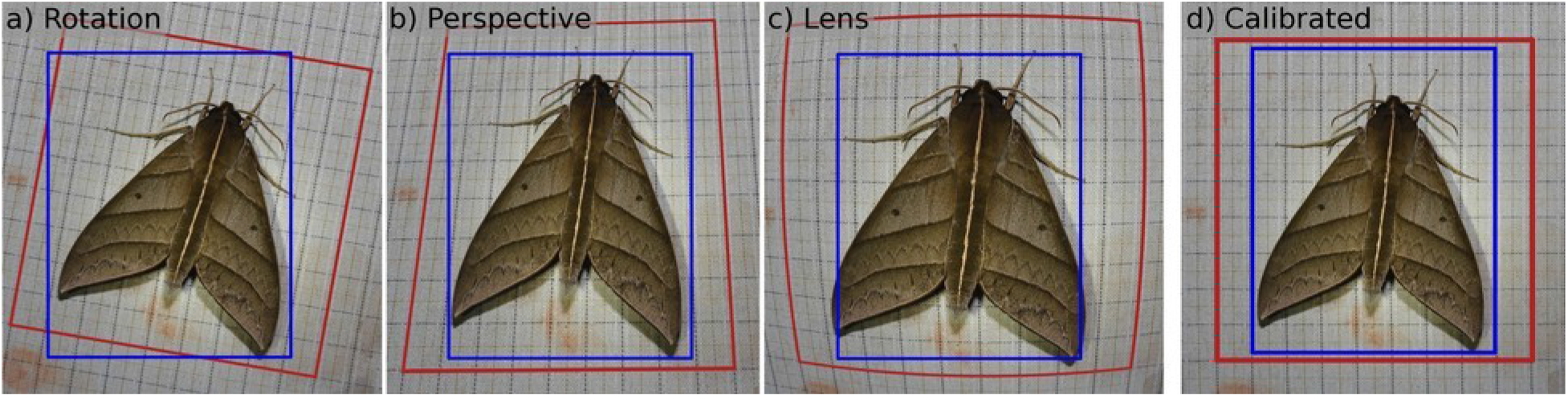
Principal distortions in a camera image: **a)** Rotation: mixing of x- and y-axes **b)** Perspective: pixel scale gradient across the image and **c)** Lens/Curvilinear: curving of straight lines. The procedure described here results in the **d)** calibrated image, in which both shape and scale distortions are corrected. The blue and the red rectangles, which represent the orthogonal X-Y axes of the image coordinates and of the subject, respectively, should become parallel to each other in the calibrated image.

**Figure 3.**
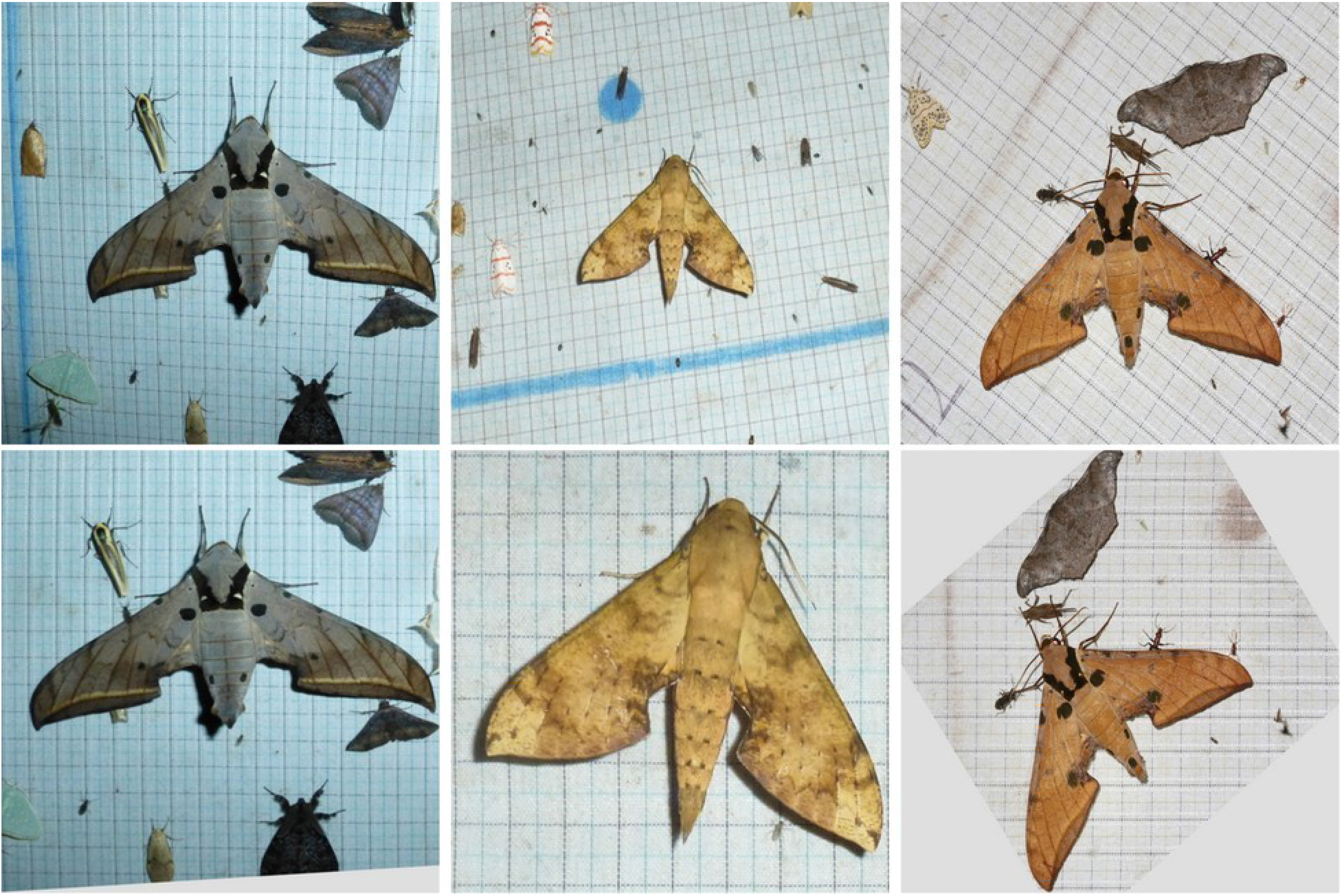
**Top row:** Examples of distorted raw images. **Bottom row:** Final de-distorted/calibrated images.

### B. Image Processing

The post-imaging calibration procedure consists of 3 tasks including de-distortion, error estimation and trait measurement, all of which require the identification of image landmarks. We developed 3 functions in R (version 3.4 or higher; R Core Team, 2013) using the libraries ***imager*** (v. 0.41.2 or higher, Bache & Wickham, 2007), ***magick*** (v. 2.0 or higher, Ooms, 2018), ***jpeg*** (v. 0.1.8 or higher, Urbanek, 2014), ***reshape2*** (v. 1.4.3 or higher, Wickham, 2007) and ***plyr*** (v. 1.8.4 or higher, Wickham, 2011); and the image processing software suite ***ImageMagick*** (v. 6.9 or higher; ImageMagick Studio, L. L. C., 2008). The 3 functions are used to (i) output the pixel coordinates of salient image landmarks, (ii) create and execute command files for dedistorting images, and (iii) estimate error heuristics to determine the quality of the final image. Manual intervention was largely limited to locating the landmarks, which took less than a minute per image.

The tasks were successfully executed under Linux Ubuntu (18.04), Windows and Mac operating systems.

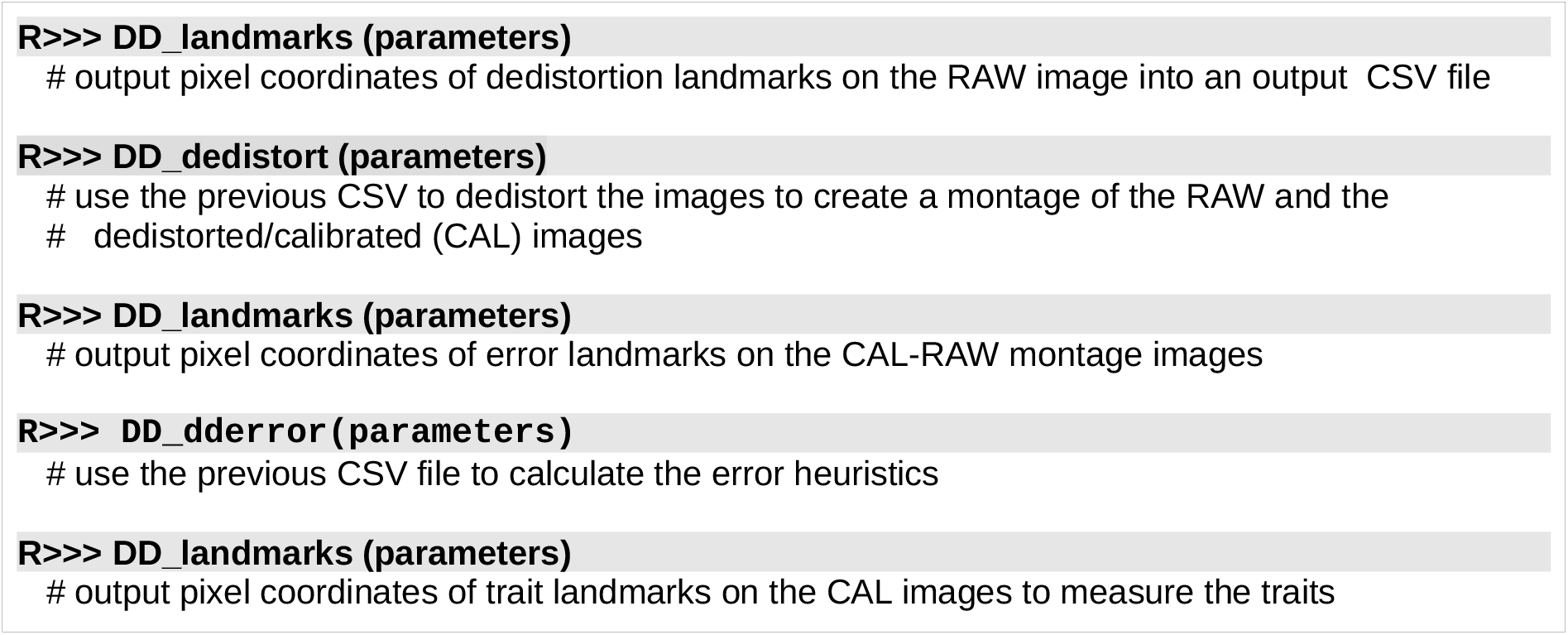

#### i) Landmarks on the Image

The function ***DD_landmarks*** outputs a user-specified number of landmarks in a CSV^1^ file. On execution it launches the selected graphic pane (X11, Quartz, or Windows) and displays all the images in the folder with the specified file tag by turn (Figure 4a).

**Figure 4:**
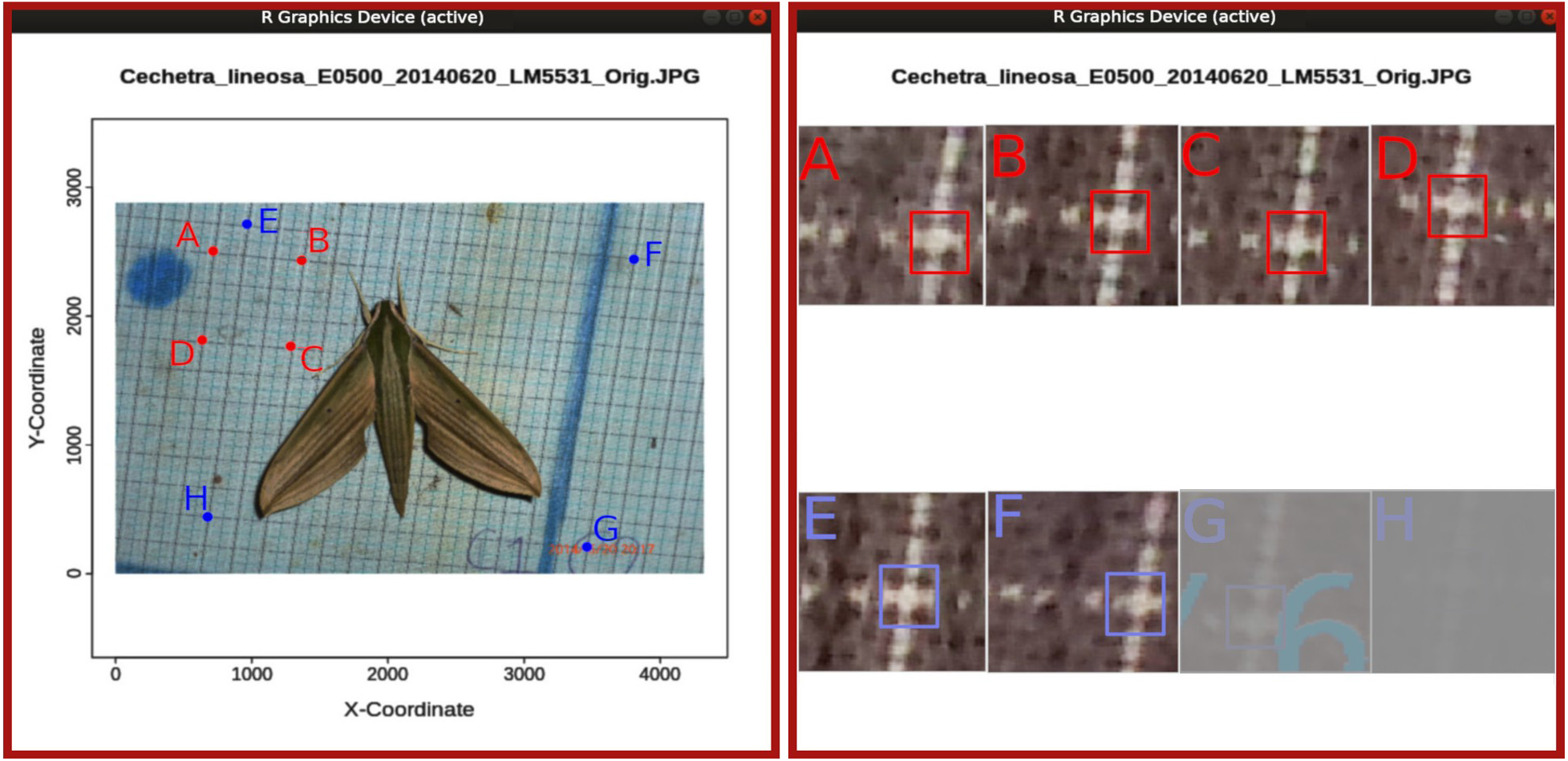
The Graphical User Interface (GUI) of the two-step Landmark function. **Left:** First, the image is shown in full for approximately marking the pixels, e.g. A-H, using mouse clicks. **Right:** Subsequently, high contrast, magnified 100 x 100 pixel postage stamps around the marked locations are displayed in sequence for more accurate selection of the pixels.The panels suggest that the postage stamps A-F have been displayed and marked, and G and H are to follow.

The advantage of our landmark procedure over previous implementations is that pixel location is a 2-step process. In the first step the required number of landmarks are approximately located on the full image. Subsequently, the function displays a sequential montage of postage stamps of the zoomed-in, higher contrast 100×100 pixel views of the neighbourhood of the landmarks (Figure 4). The user can now mark the required pixel with greater accuracy.

#### ii) De-distortion

Essentially, this task requires the user to mark two rectangles, ABCD and EFGH, using ***DD_landmarks*** as per the pattern in Figure 5. ABCD should be a *grid-corner rectangle* – i.e. its vertices lie on grid intersections – and it provides both the scale and the shape reference since the basic grid is a rectangle of known size (X: 7.0 mm, and Y: 7.3 mm in our case). The size of ABCD can be fixed in two ways:

1. Fix the same at the start (parameters ***markcode = “DD8”, ngridX***, ***ngridY***): Mark ABCD such that AB = CD = ngridX, and BC = DA = ngridY in units of the grid.
2. Mark the size on the fly (parameters ***markcode = “DD12”, ngridX***, ***ngridY***): Mark ABCD of any size, and additionally mark line segments IJ = ngridX parallel to AB, and KL = ngridY parallel to BC. The ratios of AB/IJ and BC/KL in pixels yield the size of ABCD.

**Figure 5.**
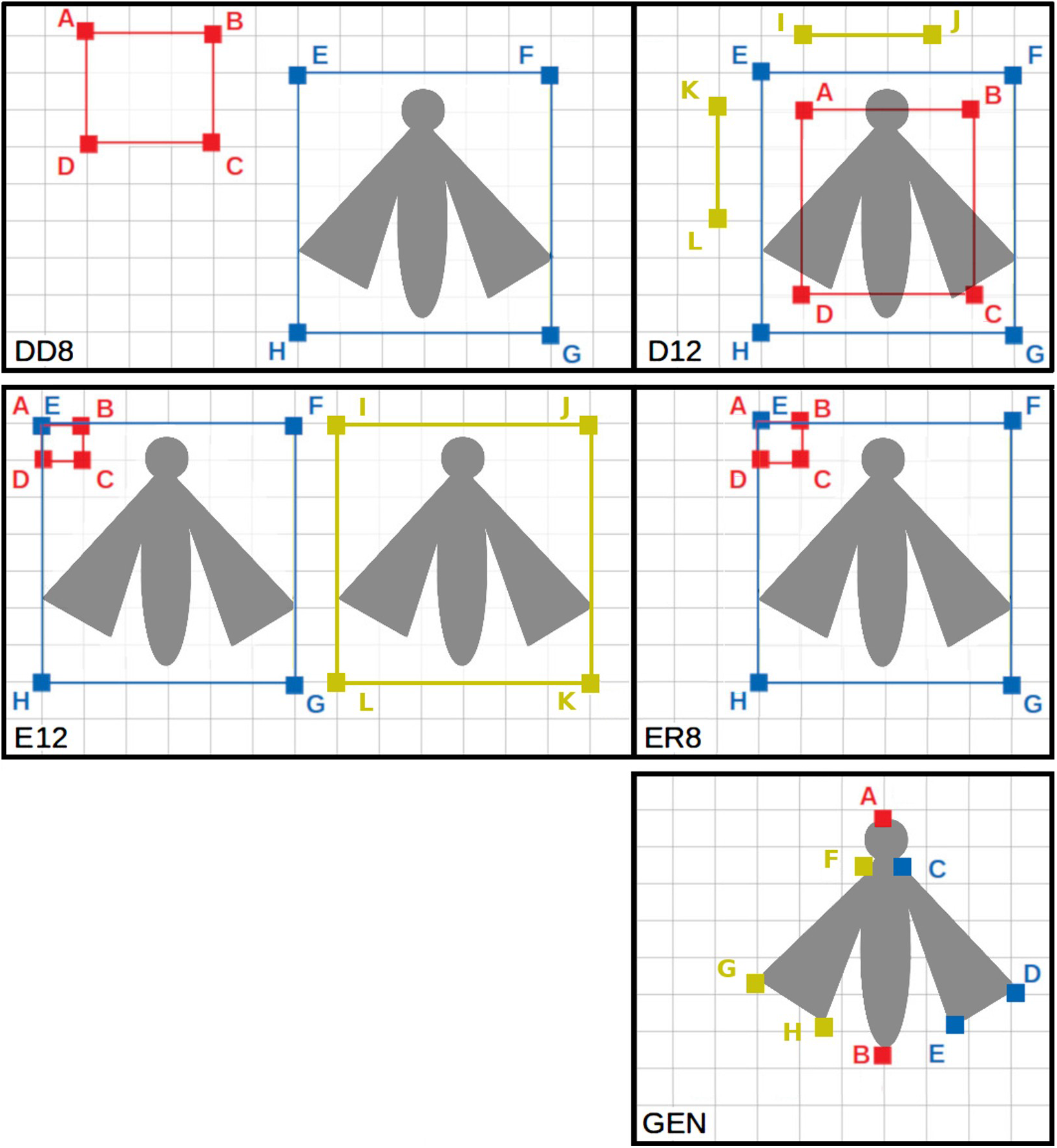
Landmark patterns for **(a)** DD8: 8 pixel de-distortion, **(b)** DD12 pixel dedistortion, (c) ER8: 8 pixel dedistortion error estimate, and (d) ER12:12 pixel dedistortion error estimate. The rectangle ABCD, and the points I-J-K-L are always marked on grid intersections, as are EFGH in ER8 and ER12. There is also the general option “GEN” for a user-specified number of landmarks (e.g. for traits).

The rectangle EFGH is used to identify the region of interest on the image in both cases. Using the recorded location of ABCD on the raw image, the function ***DD_dedistortion*** (parameters: ***mmperXgrid = 7.0 mm, mmperYgrid = 7.3 mm, pixpermm = 10***) calculates transformed location of ABCD on the calibrated image (requiring that all the angles be 90°, and fixing the scale of the output image using

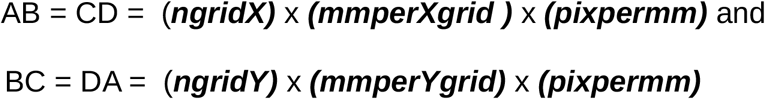

The coordinates of ABCD on the raw and the calibrated images are used as parameters of the *ImageMagick* command to dedistort the image. Currently, the ***magick*** library in R does not provide the dedistortion utility. So we create a system command/batch file which can be executed either manually (***fg_imagemagick = FALSE***) or by the R finction itself (***fg_imagemagick = TRUE***)

#### iii) De-distortion error estimation

The function ***DD_dderror*** calculates the errors in the calibration process when provided landmarks from ***DD_landmarks*** using one of the two options below:

1. parameters ***markcode = “ER8”, ngridX, ngridY)***

∘ On the calibrated image: mark the grid-corner rectangle ABCD with AB = CD = ***ngridX*** and BC = DA = ***ngridY*** as shown in Figure 5. EFGH can be any grid corner rectangle which approximately encompasses the subject. The location of A and E should coincide.
2. parameters ***markcode = “ER12”, ngridX, ngridY)***

∘ On the calibrated image: mark ABCD and EFGH as for ***ER8***
∘ On the raw image: mark grid-corner rectangle IJKL at the locations (approximately) corresponding to EFGH.

The task calculates the following errors (see Figure 2)

1. Pixel location error: Length AE is a measure of human error in marking the desired pixel, since A and E are meant to be marked on the same pixel.
2. Grid misalignment error: Segments EF and GH are meant to be horizontal, and FG and HE vertical, and their deviations from the same are a measure of the errors in the dedistortion process. A deviation θ results in a fractional length error (δL/L) = (1 – cosθ), i.e. a 5.°3 inclination of a grid line will cause a linear error of ~1%. Rotation in itself causes no change in lengths, though it can introduce errors if the scale distortions along the X and Y axes are not identical.
3. Scale error: This is the deviation from the expected scale ***pixpermm*** (= 0.1 mm per pixel for us) set in the previous task. It is measured by counting the number of pixels between a known number of grid intersections on the calibrated image. If the measured length of AB and EF in pixels are AB_P_ and EF_P_, the number of grids in EF ≡ EF_G_ = round(EF_P_/AB_P_). The expected number of pixels in EF_P-EXP_ = EF_G_ x ***mmperXgrid*** x ***pixpermm***. The percentage scale error δS_EF_ = 100 x (EF_P_ – EF_P-EXP_)/EF_P-EXP_. A similar scale error can be calculated for the other 3 sides.
4. Perspective distortion: Perspective changes a rectangle into a trapezium, and results in a gradient in scale across the image. It occurs when the plane of the camera is not parallel to the plane (of the subject) in which the measurements are to be made. The quantity below, which is zero for a rotated rectangle or a parallelogram (shear distortion) and non-zero for a trapezium, provides a heuristic estimate of the perspective error:

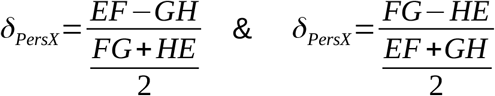 One can also estimate the efficacy of the procedure by comparing the perspective error before and after de-distortion. We combined the values for the above 4 errors in the X- and Y-direction to derive a standard deviation for the distribution of the entire sample of moths as an estimate of the accuracy of the procedure.
5. Any gap between the plane of the insect wings and the gridded cloth reference, even when they are parallel, would over-estimate lengths, since the wings, which are nearer, are projected against the farther screen. The fractional error due to this gap should be equal to the ratio of the wing-screen and camera-wing distances.

Other distortions (e.g. shear, pin-cushion, etc) can be quantified individually but are not necessary for our purpose. Essentially, the scale error is much larger than the others and we suspect all the others are reflected in it. Images with error values much larger than expected from the distribution for the whole sample were re-processed to eliminate human error. Those which continued to have high errors were removed from subsequent analyses.

#### iv) Measuring traits on the de-distorted image

We measured the following traits on the de-distorted images using the landmark pattern A-B … C-D-E … F-G-H shown in Figure 5:

- Body-length = AB (length of the segment AB)
- Thorax-width = CF
- Wing triangles ≡ C-D-E and F-G-H. Using wing terminology, CD and FG are the (right and left) costa, DE and GH are the termen, and EC and HF are the dorsa.
- Body-volume, modelled as a bi-cone (spindle-shaped)

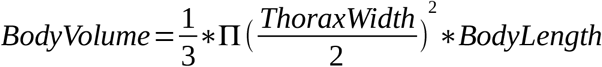
- Wing area, calculated from the lengths of the sides

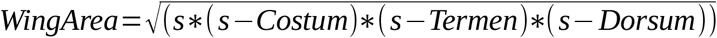

where,

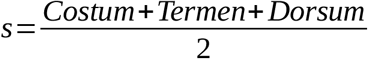
- 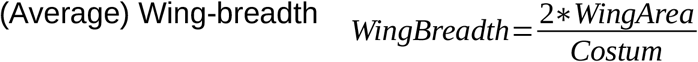

#### v) Curation of trait values

We identified outliers in the distribution of traits (body-length, body-volume, wing-length = costum, and wing-area), either separately for each taxon or for the entire community, as appropriate, in terms of the deviation from the mean in units of the standard deviation. Species with substantial sexual dimorphism should result in a bimodal distribution but we did not encounter any such in our data.

Wings held at an angle to the gridded screen will acquire distortion when the background grid is de-distorted (Figure 6). This could result in a large difference between the trait values of the left and the right wings. We used the normalised difference between the left and the right wing dimensions to identify poorly positioned moths. Normalising this difference allowed us to combine the data from all species into a single distribution for better estimation of the standard deviation.

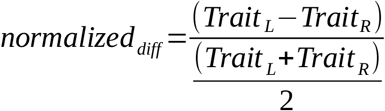

**Figure 6.**
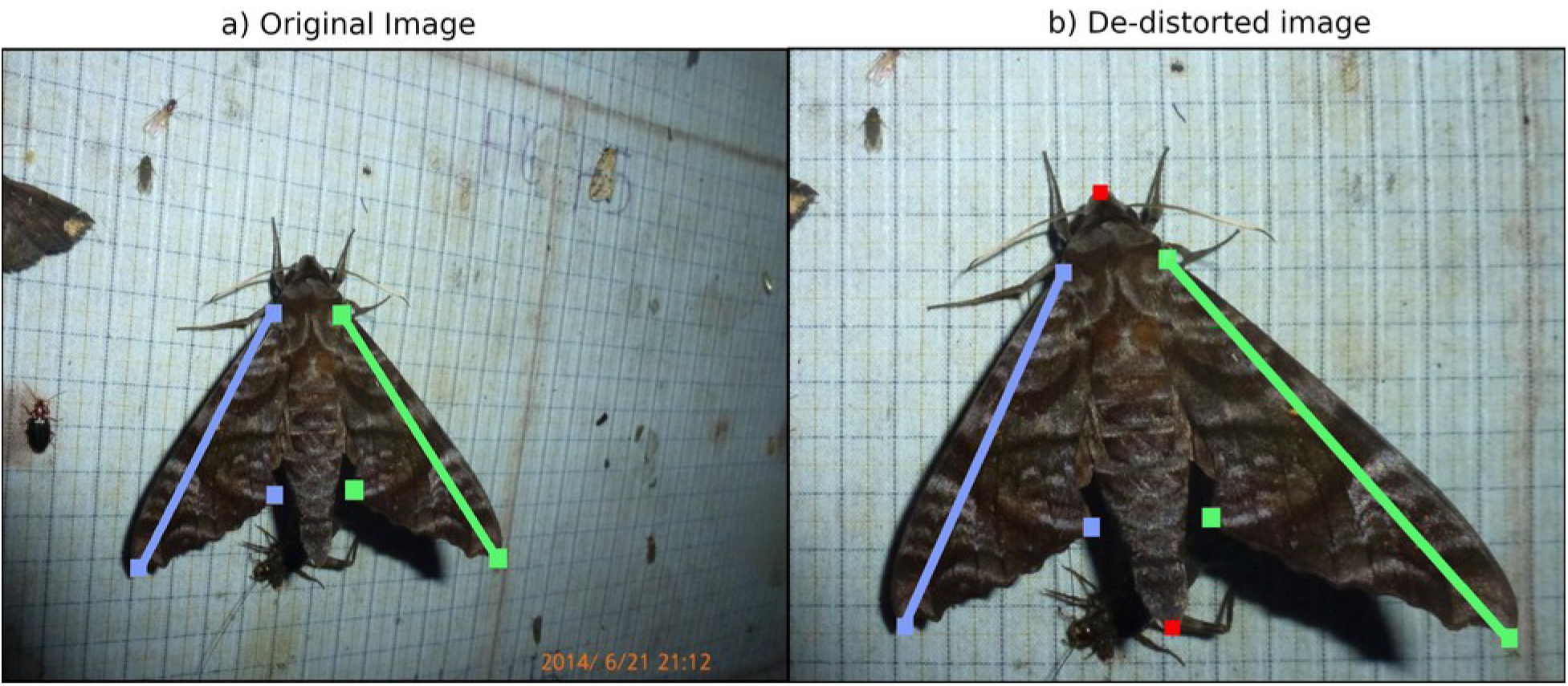
A left-right asymmetry is introduced by the de-distortion procedure when the wing is not parallel to the screen – the shadow under the right wing suggests a large gap. The original image shows a well-positioned moth with symmetrical wings on a screen which is angling away. De-distortion transformed the distorted grids into rectangles and imposed the same, but inappropriate, correction on the subject.

Since we were interested in the relationship between the body-volume and wing-area we also used the residuals from their linear regression to identify and flag outliers. Any pair of related traits of interest may be used in a similar manner. We used the following transformed values for the regression

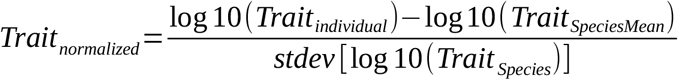

#### vi) Comparison with specimens measured using calipers

We assessed the accuracy of the method by comparing body lengths from photogrammetric and direct (using a scale of least count 1 mm) measurements for 105 specimens that had been both imaged and collected.

## Results

The linear regression between the body lengths from specimens and images, for the 105 individuals with both measurements, is shown in Figure 7 (y-intercept = 1.71 ± 1.10, slope = 0.96 ± 0.02, R^2^ = 0.94). The value is consistent with zero y-intercept at 1.6 σ. A ratio test of the two, equivalent to forcing a linear regression with y-intercept = 0, yielded a 95% confidence interval for the mean of [0.996, 1.010] against the expected value 1.0 (Table 1), which argues for no bias. Similarly, a difference test yielded 95% C.I. for the mean of [-0.22, +0.41] mm against the expected value 0.0.

**Figure 7.**
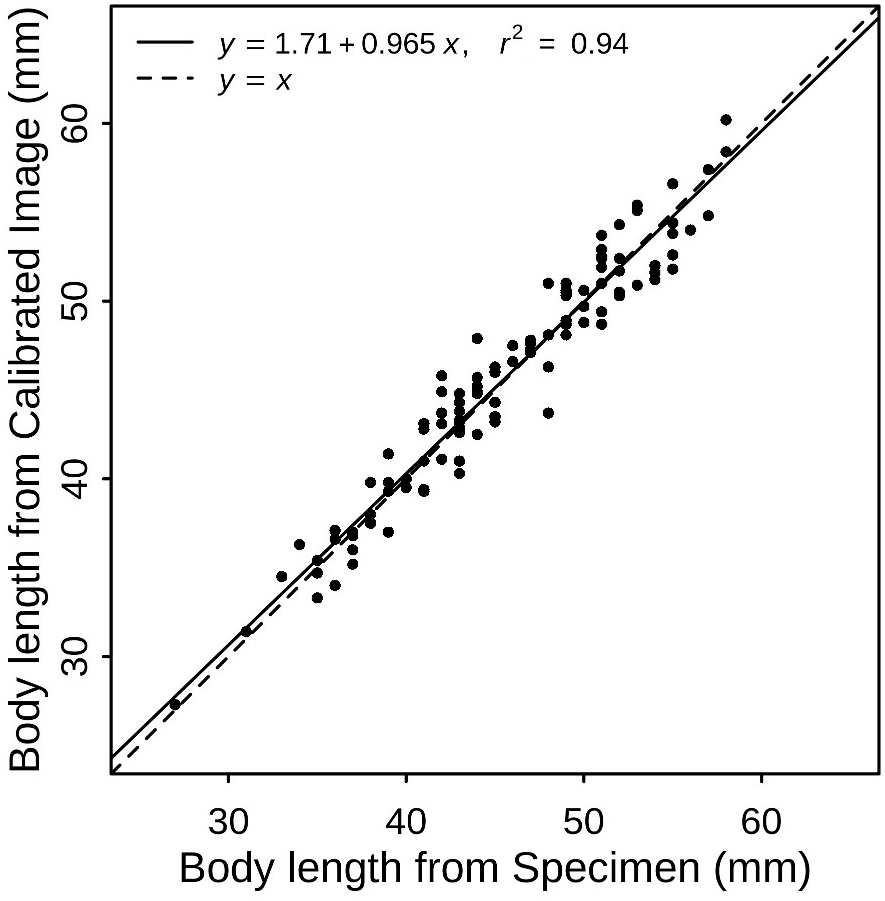
Comparison of body lengths from photogrammetry and direct specimen measurement using a scale.

**Table 1.**
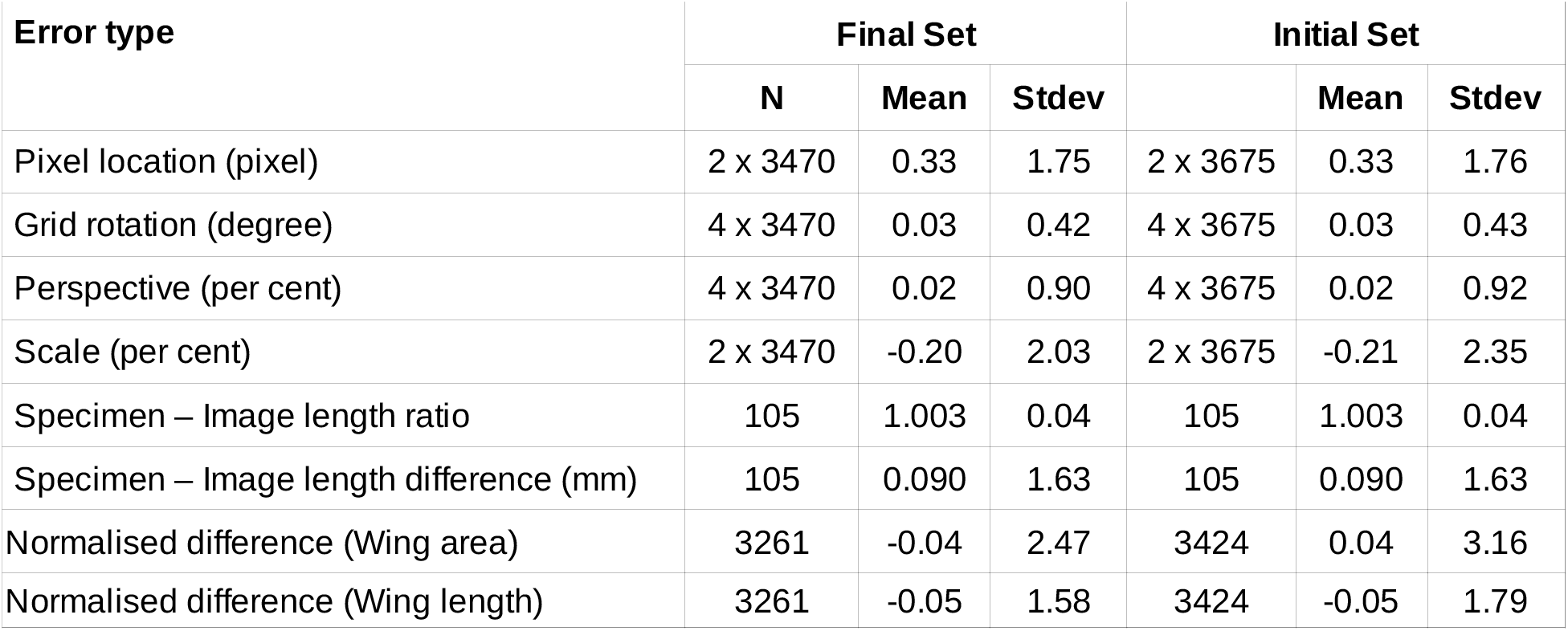
Statistics of the error metrics. Each image may yield 1, 2, or 4 independent values of the metric. Robust statistics have been calculated using 3σ clipping.

| Error type | Final Set |  |  | Initial Set |  |  |
| --- | --- | --- | --- | --- | --- | --- |
|  | N | Mean | Stdev |  | Mean | Stdev |
| Pixel location (pixel) | 2 x 3470 | 0.33 | 1.75 | 2 x 3675 | 0.33 | 1.76 |
| Grid rotation (degree) | 4 x 3470 | 0.03 | 0.42 | 4 x 3675 | 0.03 | 0.43 |
| Perspective (per cent) | 4 x 3470 | 0.02 | 0.90 | 4 x 3675 | 0.02 | 0.92 |
| Scale (per cent) | 2 x 3470 | -0.20 | 2.03 | 2 x 3675 | -0.21 | 2.35 |
| Specimen – Image length ratio | 105 | 1.003 | 0.04 | 105 | 1.003 | 0.04 |
| Specimen – Image length difference (mm) | 105 | 0.090 | 1.63 | 105 | 0.090 | 1.63 |
| Normalised difference (Wing area) | 3261 | -0.04 | 2.47 | 3424 | 0.04 | 3.16 |
| Normalised difference (Wing length) | 3261 | -0.05 | 1.58 | 3424 | -0.05 | 1.79 |

We obtained images and collected 2 middle legs for DNA analysis from 4808 hawkmoth individuals, of which 3675 images had the gridded screen as background. Some of the others had rested on the surrounding vegetation or the tetrapod, while the rest could not be photographed properly due to heavy rains. These images were used to identify the individuals to (morpho)-species and de-distorted in batch mode to measure the traits. Approximately 500 images could be dedistorted during an 8-hour session.

Since our images were taken from a distance of about 30 cm, and the resting wings are mostly held flush against the screen, or at most a few millimeter away, the error due to the gap between the wing and the screen should be about 1 % (ratio of the two distances).

The statistics of the various metrics are listed in Table 1.

The distribution of perspective errors after de-distortion is shown in Figure 8^2^. The second panel in Figure 8 shows the distribution of the difference in the absolute values of the perspective error before and after the procedure. As expected, the values are mostly positive, i.e. the errors reduced, and the numbers fall sharply below zero.

**Figure 8.**
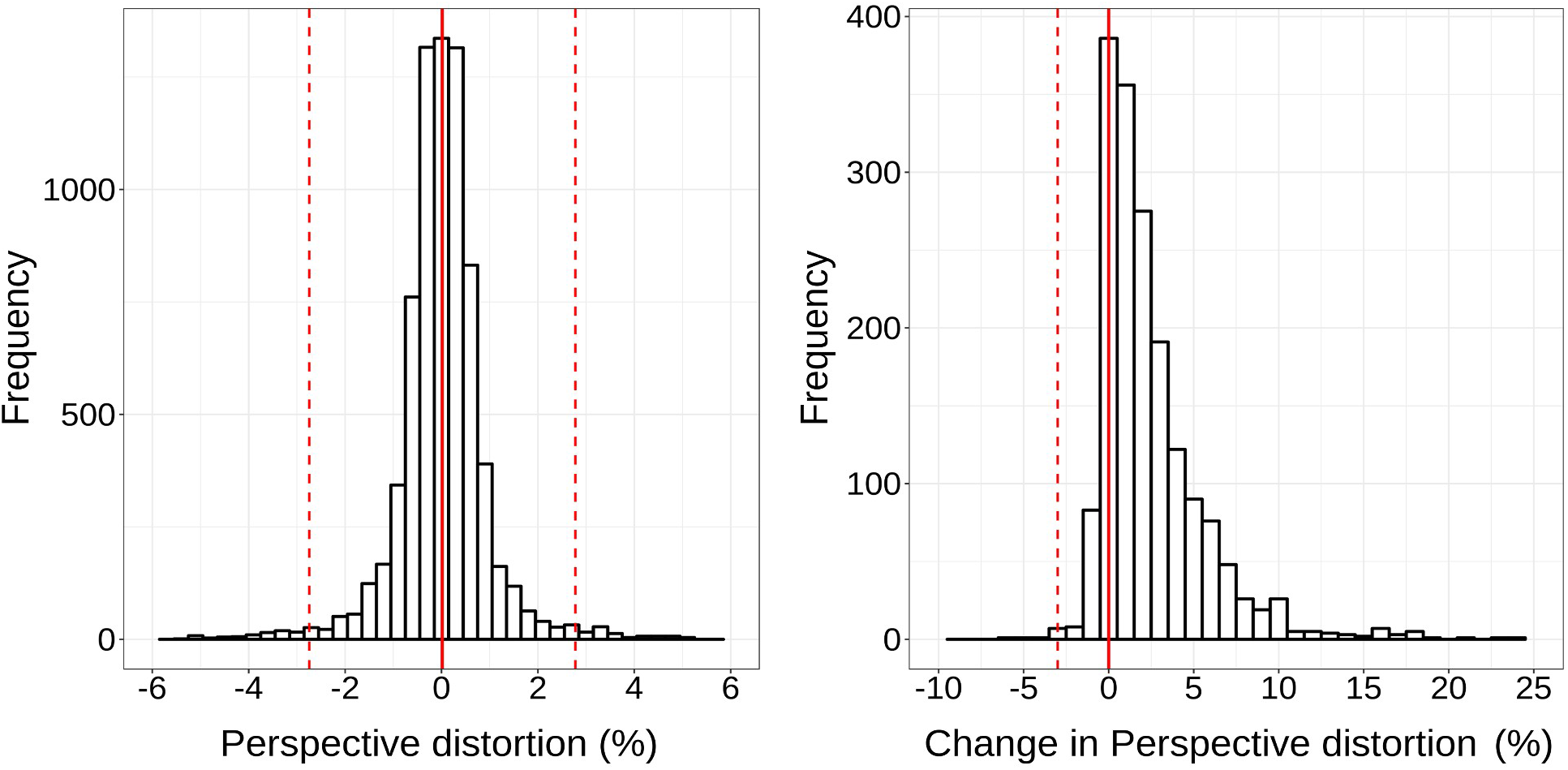
Distributions of **(Left)** perspective error in the calibrated image and (**Right**) difference between the absolute values of the perspective error in the raw and calibrated images. Note the sharp fall in numbers to the left of zero; i.e. the errors were much larger in the raw image

The distributions of the other three error metrics are shown in Figure 9. The pixel location error SD = 1.76 pixels (= 0.18 mm) corresponds to a fractional length error of 0.35% for a typical hawkmoth body length of 50 mm. The mean axis misalignment SD = 0.°43, correspond to a negligible fractional length error of 0.02%. The scale factor showed SD = 2.3%.

**Figure 9.**
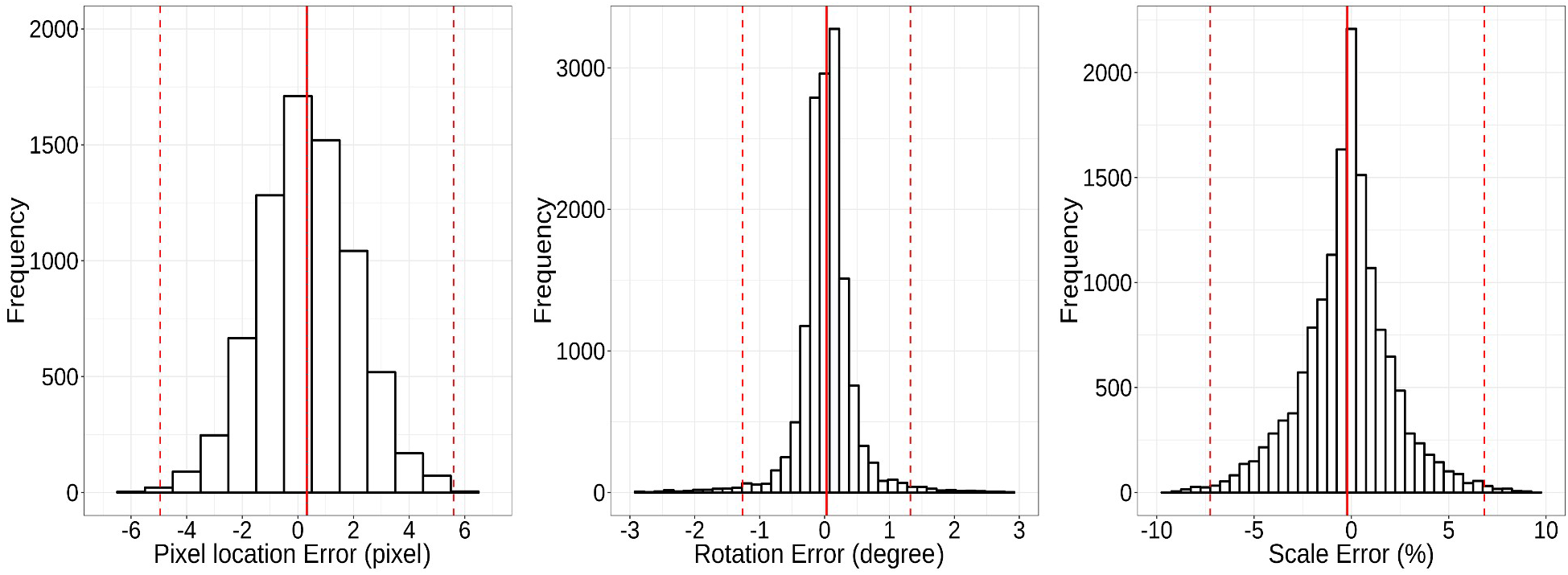
Distributions of dedistortion error-metrics: (**Left)** Error in marking the target pixel, **(Mid)** Misalignment of the grid lines from the vertical and the horizontal, and **(Right)** Scale error, i.e. the deviation from the expected number of pixels along a grid segment

The distribution for the normalized difference between left and right wing-area and wing-costum length are shown in Figure 10. The means of both the distributions are close to zero and SD = 1.6% for wing length, and 2.5% for area. We found that in some cases even though the left-right asymmetry was stark, averaging the two (as is appropriate for an individual) resulted in a value within the species distribution; presumably this effect impacts the two wings in opposite directions without appreciably changing their mean.

**Figure 10.**
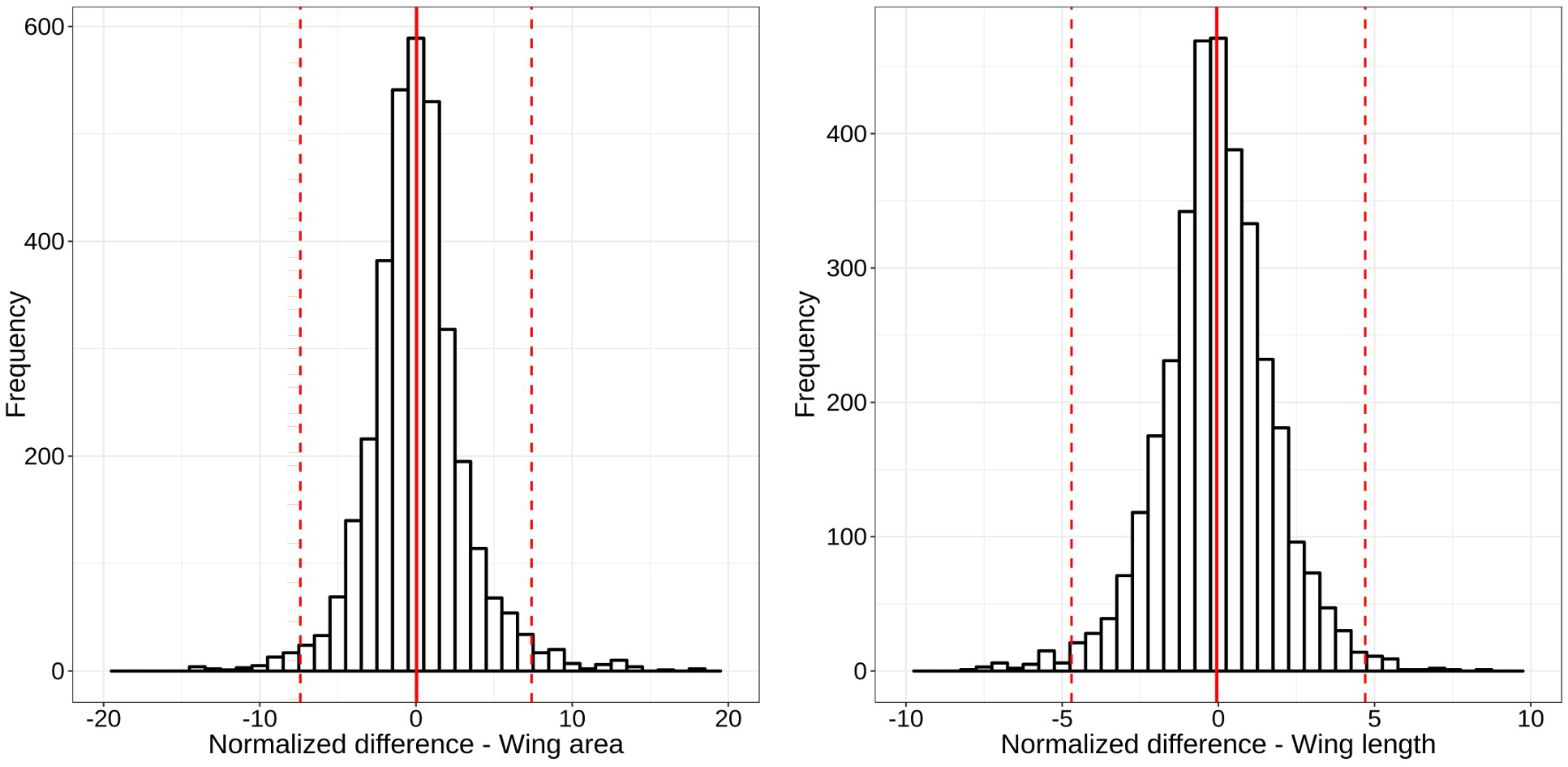
Distributions of the normalized difference between the left and right wings: **(Left)** area, and **(Right)** costum length. The difference is a measure of non-coplanarity between the wing and the screen.

We generated species-wise trait histograms for body volume and wing area, and regression plots for wing area on body volume. Figures 11 and 12 show the distributions for a single species, *Cechetra lineosa*, as an illustration.

**Figure 11.**
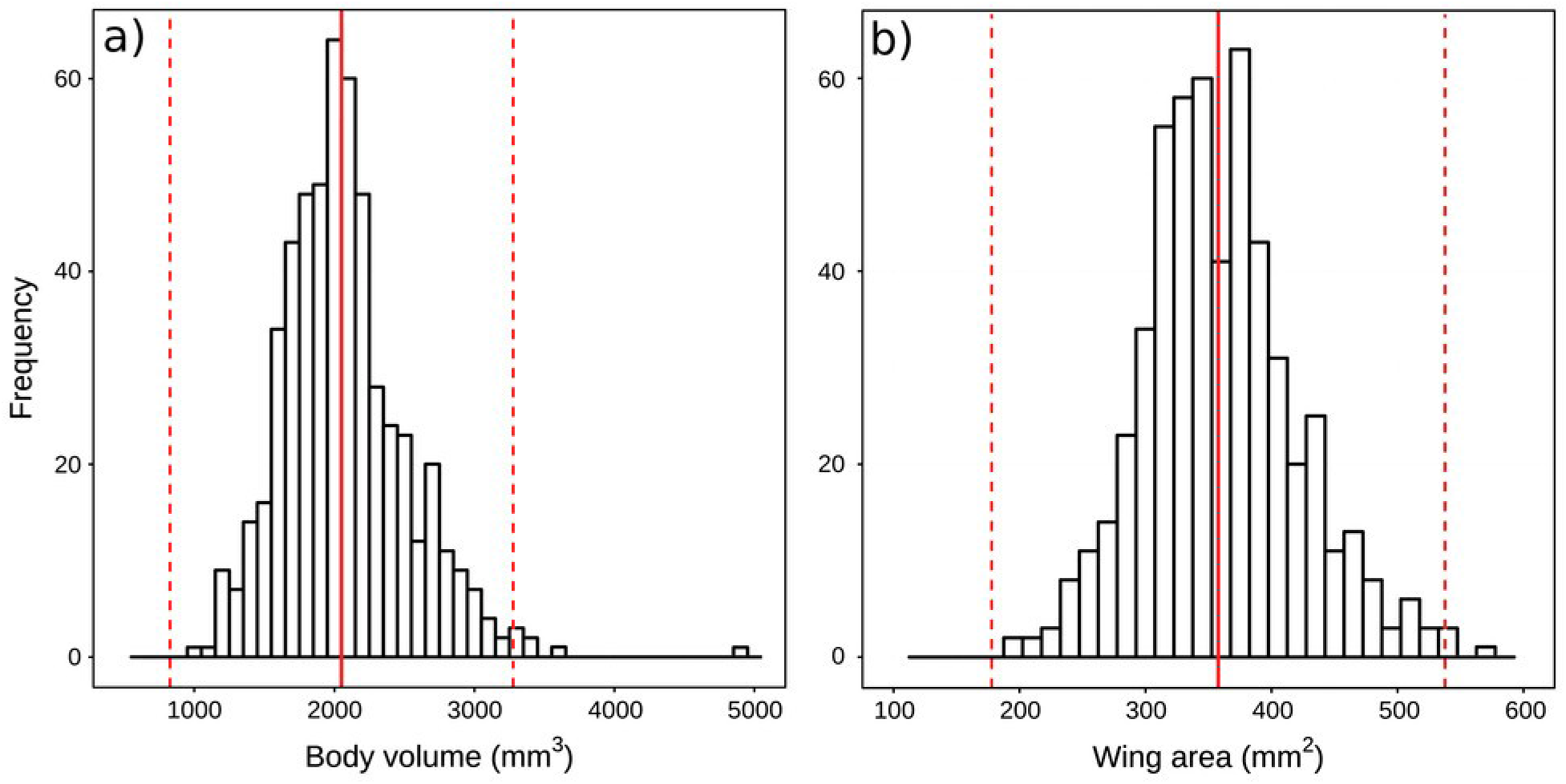
Distributions of body volume **(Left)** and wing area **(Right)** for individuals of the species Cechetra lineosa. The red lines represent the means and the 3σ values.

**Figure 12.**
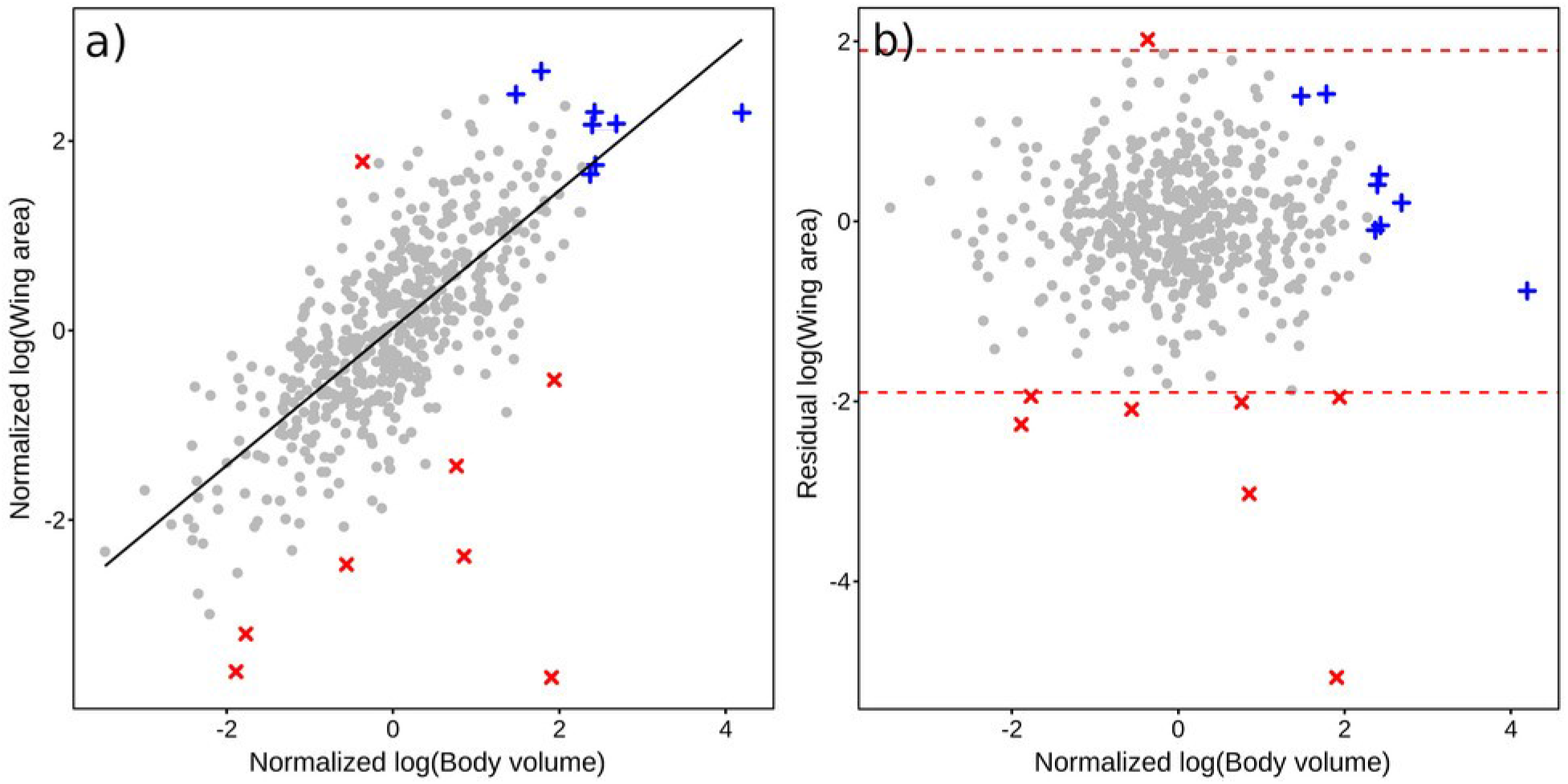
Linear regression of wing-area on body-volume **(Left)** and its residual **(Right)** for Cechetra lineosa. Regression outliers are in red and insects with only one wing measurements are in blue. Wingarea and body-volume are expressed as deviation from the species mean in units of the standard deviation.

One can define thresholds for selecting “usable” images in several ways. It can be on the basis of deviation from the mean in units of the standard deviation (e.g. reject images whose error estimate was more than 3 x SD from the mean). On the other hand when the errors are very low (e.g. only 0.01% for misalignment error), it is wasteful to eliminate images using this criterion. Alternatively, the accuracy required by the investigation can set the threshold (e.g. 10% fractional length error).

In the sample of 3675 images presented here only 14 images (0.4%) had scale errors in excess of 10% and 3470 images (94.4%) of the images had an error less than 6.1 % (3σ after robust clipping).

## Discussion

We have demonstrated a simple, inexpensive method for obtaining robust morphometric measurements from digital images. Images of free-ranging moths on a light screen provided basic morphometrics without having to allocate considerable resources for the collection, processing and storage of specimens. This method can be applied to other phototropic insect taxa as well (e.g. *Coleoptera*, *Aphidina*, *Diptera*, *Trichoptera*, *Heteroptera* and *Hymenoptera*; see van Grunsven et al., 2014, for attracting phototropic insects) and any visible morphological trait such as body length, thorax width, head width, wing sizes, antennae length etc. The equipment and material necessary for this are all consumer-level, off-the-shelf items which makes the procedure accessible to researchers all across the world, and yet provides measurements accurate to a few per cent for tens of thousands of individuals.

Photogrammetry is currently limited to large mammals and birds. Previous methods of distortion correction from digital images have relied on additional information such as aerial photographs from multiple angles with multiple cameras (Gerum et al., 2017) or on a checkerboard in the field of view for edge detection and distortion correction (CCTM of MATLAB; Heikkila & Siliven, 1997). In our case, the gridded screen, needed as a resting surface for phototropic insects, provides a dedistortion reference at no extra cost.

The landmark task in R developed for this work (***DD_landmarks***) differs from previous utilities (e.g. in the R-package Geomorph, Adams & Otàrola-Castillo 2013) by employing a two-step marking procedure for better accuracy in locating pixels.

The human effort required was only about 1 minute per individual in the field, and in the lab; a person can process about 500 images a day in the lab. Both these tasks can be easily carried out by personnel with little or no academic or technical background. As importantly, the procedure yielded quantifiable errors on images which are available for repeated measurements.

The comparison of lengths from specimens and images showed that any systematic difference between the two is less than 0.04 mm (half pixel) at 95% confidence level. The overall precision, reflected in the standard deviation of the scale factor, was 2.3%, perhaps arising from the stretchable nature of the fabric and variable humidity conditions. This was adequate for our purpose, especially since intraspecific variation is much larger (e.g. Figure 11), but a less stretchable screen can reduce the error even further.

The rapid rate of environmental change, from both land-use pattern and climate changes, are expected to trigger substantial responses in many ecosystems. One can directly track these responses by monitoring changes in species communities and in their key morphological traits since individuals interact with and respond to the environment via their traits. Such multi-epoch monitoring would require the sampling of hundreds of thousands of individuals every year, and can benefit from not having to collect tens of thousands of specimens every time. Our method provides a way of accumulating large amounts of reliable trait data at minimal cost, especially in such situations. The target hawkmoths comprised less than 1% of all the moths individuals arriving at the light screen in our study. Having to allocate limited resources to collecting and processing all the moth specimens visiting the light trap would have forced us to scale down the intended study.

Even in the case of collected specimens, this technique would make expensive instruments like a calibrated microscope unnecessary except for high precision measurements of very small subjects.

This method is particularly good for nocturnal phototropic insects, and the fact that such taxa form the bulk of biodiversity makes for its wide applicability across the globe. We acknowledge that many morphometric studies may require the collection of full specimens, even in large numbers. The photogrammmetric technique presented here obviates the necessity of a large expenditure on specimen collection, processing and maintenance, where it is not essential, and yet yields reliable morphometric data which can be reexamined by others at a later date.

## Supporting information

SupportingInformation1

## ACKNOWLEDGMENTS

RA acknowledges financial support through a grant (No. SR/SO/AS66/2011) from the Department of Science and Technology, Government of India, and Nadathur Trust, Bengaluru. We thank the Forest Department of Arunachal Pradesh for their assistance and research permits (CWL/G/13(17)/06-07/Pt-III/4194-95 and CWL/G/13(95)/2011-12/Pt.II/660-62 during 2011-2015). This work would not have been possible without the enormous support of diverse kinds provided by Mr. Indi Glow, Nima Tsering and other members of the Singchung Bugun community. We thank Srikrishna Sekhar for help with an earlier version of the software.

## STATEMENT OF AUTHORSHIP

RA designed the project, MM carried out the computations and image processing; all the rest including collection of field data, analysis, and writing of this manuscript were shared by both. We declare that we have no competing interests.

## DATA ACCESSIBILITY STATEMENT

All relevant information in support of the results presented here will be archived in the recommended public repository DRYAD, upon acceptance of the manuscript.

1. We have uploaded the data file (as a Libre-office spreadsheet) for the referee.
2. We should also be happy to deposit the source code of the three R functions used in the analyses in any public repository recommended by the journal

## Footnotes

1 CSV: a text file in the Comma-separated-value format, which can be read by any spreadsheet software like MS-EXCEL or LibreOffice

2 In the plots in Figures 8-11, the solid red line represents the mean and the dotted lines on either size are at thrice the standard deviation.

